# Progerin Expression in Alveolar Epithelial Cells Exacerbates Inflammatory Responses to SARS-CoV-2 Infection

**DOI:** 10.64898/2026.11.14.766274

**Authors:** Cesar L. Moreno, Anupriya Aggarwal, Sophia Trabish, Desi Veleva, Stuart Turville, Justin Cooper-White, Ernst Wolvetang, G. Gregory Neely

**Affiliations:** The Dr. John and Anne Chong Lab for Functional Genomics, Charles Perkins Centre and School of Life & Environmental Sciences, The University of Sydney, Sydney, NSW, Australia; The Kirby Institute, University of New South Wales, Sydney, NSW, Australia; Australian Institute for Bioengineering & Nanotechnology (AIBN), The University of Queensland, St. Lucia, Brisbane, Australia; School of Chemical Engineering, The University of Queensland, Brisbane, Queensland 4072, Australia

**Keywords:** progerin, induced pluripotent stem cells, alveolar type II cells, cellular ageing, SARS-CoV-2, inflammatory response

## Abstract

Age is a major determinant of severe coronavirus disease 2019 (COVID-19), yet the cell-intrinsic consequences of ageing for SARS-CoV-2 infection remain difficult to examine in human experimental systems. Induced pluripotent stem cell (iPSC)-derived tissues provide accessible models of infection but generally lack hallmarks of ageing. Here, we employed an iPSC line containing an inducible progerin cassette and differentiated these cells towards an alveolar type II-like epithelial fate. Progerin induction in differentiated cells increased DNA-damage-associated γH2AX staining and mitochondrial reactive oxygen species, reduced the heterochromatin-associated mark H3K9me3, and induced transcriptional programs associated with extracellular matrix remodeling, fibrosis and inflammation. Following infection with authentic SARS-CoV-2, progerin-expressing cultures exhibited a greater perturbation of the host transcriptome, including increased interferon-associated and inflammatory gene signatures. Transcriptomic analysis support higher viral RNA abundance in the aged model, and this was further confirmed by functional assays that showed increased infectious activity from condition media. Together, these data establish a temporally controlled model of selected ageing-associated phenotypes in human alveolar epithelial-like cells and suggest that progerin-induced cellular stress can amplify epithelial responses to SARS-CoV-2. More broadly, these findings distinguish epithelial-intrinsic inflammatory priming from the multicellular pathology of severe COVID-19.

## Introduction

Older age is consistently associated with an increased risk of hospitalisation and death following SARS-CoV-2 infection^1,2^. Although comorbidities and age-associated changes in systemic immunity contribute to this relationship, the role of cellular ageing within the lung epithelium remains incompletely understood. Distinguishing intrinsic changes in susceptible epithelial cells from the effects of immune-cell recruitment and tissue-wide inflammation requires experimentally tractable human models in which ageing-associated features can be evaluated independently.

Alveolar type II (AT2) epithelial cells contribute to alveolar homeostasis, surfactant production and epithelial repair and are among the lung cell populations susceptible to SARS-CoV-2. Pluripotent stem cell-derived AT2-like cells reproduce features of viral infection, including epithelial injury, antiviral signalling and responses to antiviral compounds^3^. However, their developmental immaturity and the rejuvenating effects of pluripotent reprogramming constrain their use for interrogating age-associated susceptibility and disease responses. A model that separates lineage differentiation from the experimental induction of ageing-associated stress could help address this limitation.

Progerin is an aberrantly processed form of lamin A associated with Hutchinson–Gilford progeria syndrome and has previously been used to elicit ageing-associated phenotypes in iPSC-derived cell types, including neurons^4^. Progerin expression does not reproduce the full complexity of physiological ageing, but provides a controllable means of perturbing nuclear architecture and stress pathways. We therefore adapted an inducible AAVS1-targeting strategy in human iPSCs^5^ to express progerin during lung-directed differentiation after early lineage specification. Using this system, we examined cellular stress, the baseline epithelial transcriptome, host responses to live SARS-CoV-2 and an indirect measure of released viral infectivity.

## Results

### Generation of a tunable, inducible progerin iPSC model

To enable temporal control of progerin expression, we modified a previously described inducible AAVS1-targeting vector^5^ to encode a progerin–GFP fusion protein. The resulting doxycycline-responsive line, hereafter designated dPRO, incorporated the expression cassette at the AAVS1 safe-harbour locus (Fig. 1A). After clonal isolation, targeting was assessed by junction-specific genomic PCR, which produced the expected amplicon in edited cells but not in the parental control (Fig. 1B), confirming bi-allelic integration.

**Figure 1.**
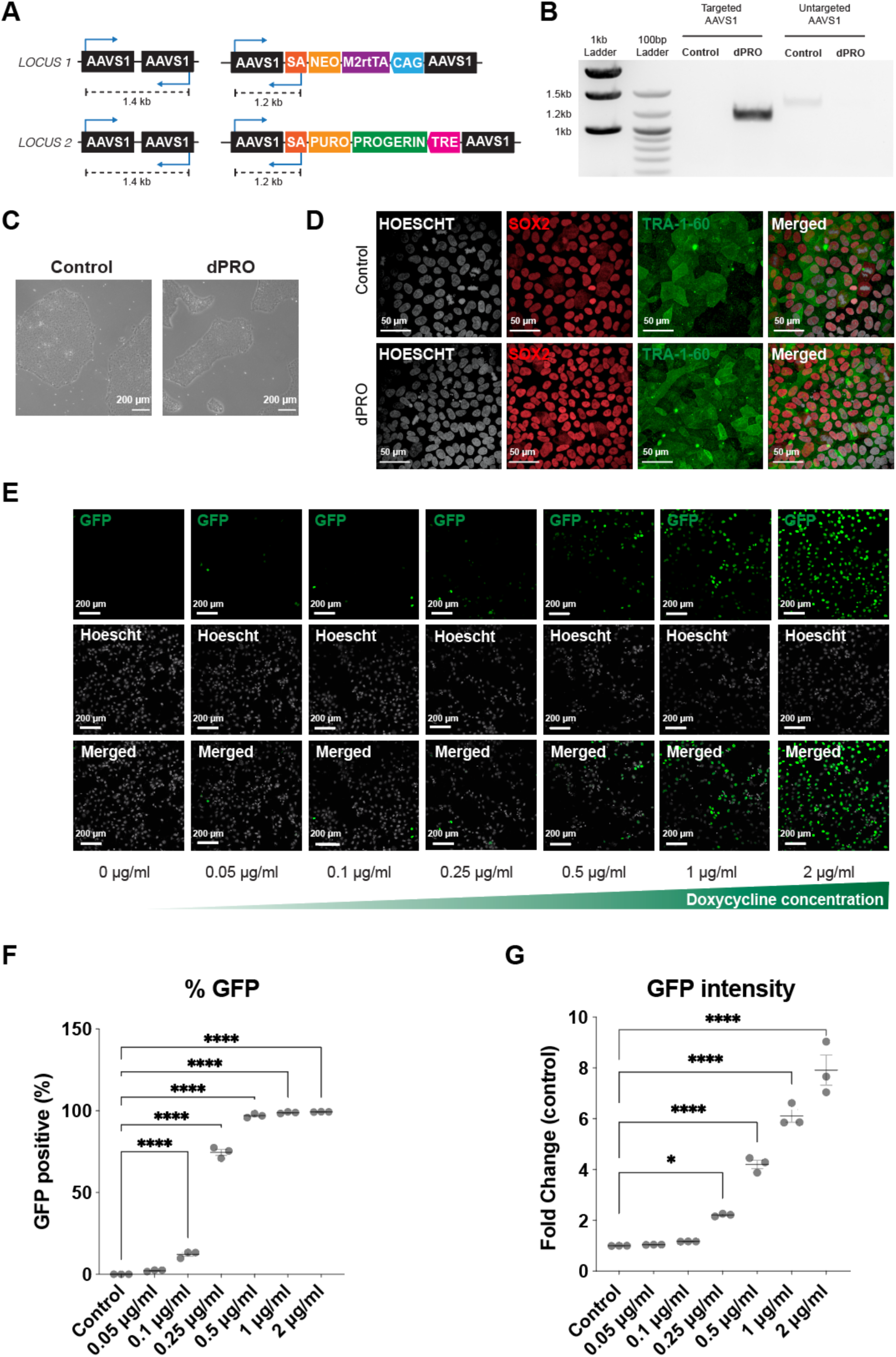
An AAVS1-targeted human iPSC line enables tunable induction of progerin. (A) Schematic of the AAVS1-targeted inducible progerin–GFP cassette and junction PCR strategy. (B) Genomic PCR products from targeted and control cells. (C) Representative colony morphology of parental and edited iPSCs. (D) Immunofluorescence of pluripotency-associated SOX2 and TRA-1-60. (E) Representative fluorescence microscopy following induction with increasing doxycycline concentrations. (F) Percentage of GFP-positive cells and (G) relative GFP fluorescence intensity after induction. N=3 experimental repeats. Error bars = SEM. Statistical significance was determined by One-Way ANOVA followed by Dunnets * P < 0.05, ****, P < 0.0001.

Edited iPSCs maintained the characteristic colony morphology of the parental line (Fig. 1C). Immunofluorescence demonstrated comparable expression of the pluripotency markers SOX2 and TRA-1-60 in edited and parental cells (Fig. 1D); OCT4 and SSEA-4 immunofluorescence further supported retention of pluripotency-associated marker expression in both parental and dPRO iPSCs (Supplementary Fig. 1A). Thus, incorporation of the inducible cassette did not compromise the pluripotent state under the culture conditions examined.

Progerin–GFP induction was then assessed across a doxycycline dose range after 24 hours (Fig. 1E–G). At 0.05 µg mL⁻¹ doxycycline, approximately 2.2% of cells were GFP-positive, whereas induction at 1–2 µg mL⁻¹ resulted in GFP positivity in more than 98% of cells. Increasing doxycycline from 1 to 2 µg mL⁻¹ further increased mean GFP fluorescence by approximately 30%, despite near-saturating induction frequencies (Fig. 1F,G). These findings establish a tunable, nearly population-wide inducible system suitable for activation after lineage specification.

### Progerin induces selected ageing-associated phenotypes in AT2-like cells

We next differentiated parental and dPRO iPSCs towards an alveolar type II-like epithelial fate using a staged protocol involving definitive endoderm, anterior foregut endoderm and subsequent lung-directed differentiation (Fig. 2A). During differentiation, pluripotency-associated *NANOG* expression declined while the lung-lineage marker *NKX2-1* increased. Expression of ACE2 and the AT2-associated marker *SFTPC* also increased relative to undifferentiated iPSCs (Fig. 2B). Bulk RNA sequencing at approximately day 23 further showed enrichment of transcripts associated with the alveolar epithelial program, including *SFTPC*, *CTSH* and *ABCA3*, accompanied by reduced expression of pluripotency-associated genes (Fig. 2C). A complementary heatmap of RNA-sequencing profiles showed distinct gene-expression patterns in undifferentiated iPSCs and day-23 AT2-like cultures, with the two sample groups separating by hierarchical clustering (Supplementary Fig. 1B). The heatmap encompassing differentially expressed genes (adjusted p value < 0.05) displays relative expression as gene-wise Z-scores. The differentiated cultures are therefore referred to as AT2-like cells rather than fully mature primary AT2 cells.

**Figure 2.**
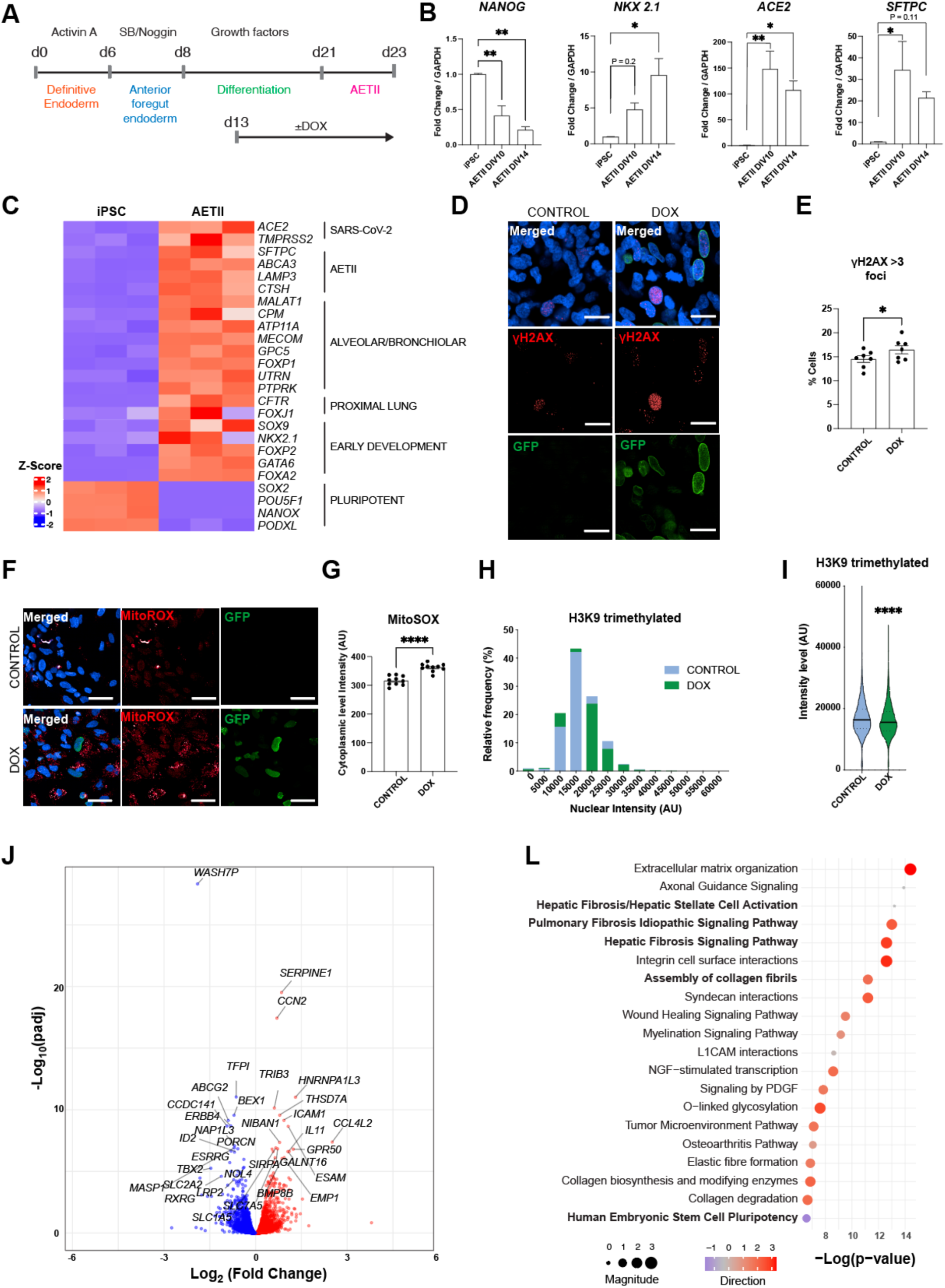
Progerin induces selected ageing-associated phenotypes in stem cell-derived AT2-like cells. (A) Differentiation and progerin-induction schedule. (B) RT-qPCR analysis of NANOG, NKX2-1, ACE2 and SFTPC during differentiation. N=3 experimental repeats. Statistical significance was determined by One-Way ANOVA followed by Dunnets * P < 0.05, **, P < 0.01. (C) Heatmap of selected lineage-associated transcript levels in iPSCs and differentiated AT2-like cells. N=3 experimental repeats. (D,E) Representative γH2AX immunostaining and quantification. Statistical significance determined by Students t-test, * P < 0.05. N=7 experimental repeats. (F,G) MitoSOX staining and quantification. Statistical significance determined by Students t-test, * P < 0.05. N=7 experimental repeats. (H,I) Distribution and summary of nuclear H3K9me3 signal. N=7 experimental repeats. (J) Differentially expressed genes following progerin induction. Statistical significance determined by Students t-test, * P < 0.05. N=3 experimental repeats. (K) Selected IPA pathways associated with the progerin response.

Doxycycline (1μg/ml) was introduced during differentiation, with progerin–GFP assessed after 10 days of induction (Fig. 2A). Compared with uninduced controls, progerin-expressing cultures showed increased nuclear γH2AX foci (Fig. 2D,E). Cell populations in the overall population showed approximately a 15% increment in foci, and this was more prevalent (∼46%) among GFP-positive cells, supporting that the increase in nuclear damage markers is driven by progerin expression. The phenotype was not observed in undifferentiated iPSCs in the original comparison (Supplementary Fig. 2C,D).

MitoSOX staining likewise indicated increased mitochondrial oxidant-associated fluorescence following progerin induction (Fig. 2F,G). In addition, nuclear H3K9 trimethylation (H3K9me3), a heterochromatin-associated histone modification, was reduced in doxycycline-treated cells (Fig. 2H,I). A senescence-associated β-galactosidase assay did not show a detectable change under the conditions tested (data not shown). Collectively, the results indicate induction of several ageing-associated stress phenotypes, without establishing a complete senescent state.

Transcriptomic profiling of induced and uninduced AT2-like cells identified 246 upregulated and 243 downregulated genes (at adjusted p-value < 0.05) (Fig. 2J). Among the transcriptional changes were increased *SERPINE1*, a mediator associated with AT2-cell senescence and pulmonary fibrosis^6^, and *CCN2*, a regulator implicated in fibrotic remodeling^7^. *TBX2*, which has been linked to senescence bypass in a distinct cancer-cell context, was reduced^8^. The inflammatory-associated transcripts *ICAM1*, a cell-adhesion molecule expressed by inflamed pulmonary epithelium^9^, *IL11* and *CCL4L2* were increased. Ingenuity Pathway Analysis (IPA) highlighted extracellular matrix organisation, wound-repair and pulmonary fibrosis-associated program (Fig. 2K). These changes suggest that progerin shifts the baseline state of differentiated alveolar epithelial-like cells towards a stress-associated, inflammatory and profibrotic transcriptional program.

### SARS-CoV-2 infection elicits stronger inflammatory transcriptional responses in progerin-expressing cultures

To determine whether progerin induction altered the epithelial response to SARS-CoV-2, differentiated cells were exposed to authentic virus following the induction interval (Fig. 3A). Flow cytometry confirmed that differentiated AT2-like cells expressed surface ACE2 more prominently than undifferentiated iPSCs (Fig. 3B). Virus exposure produced cytopathic effects and detectable SARS-CoV-2 antigen staining in differentiated cultures (Fig. 3C,D), supporting their permissiveness to infection.

**Figure 3.**
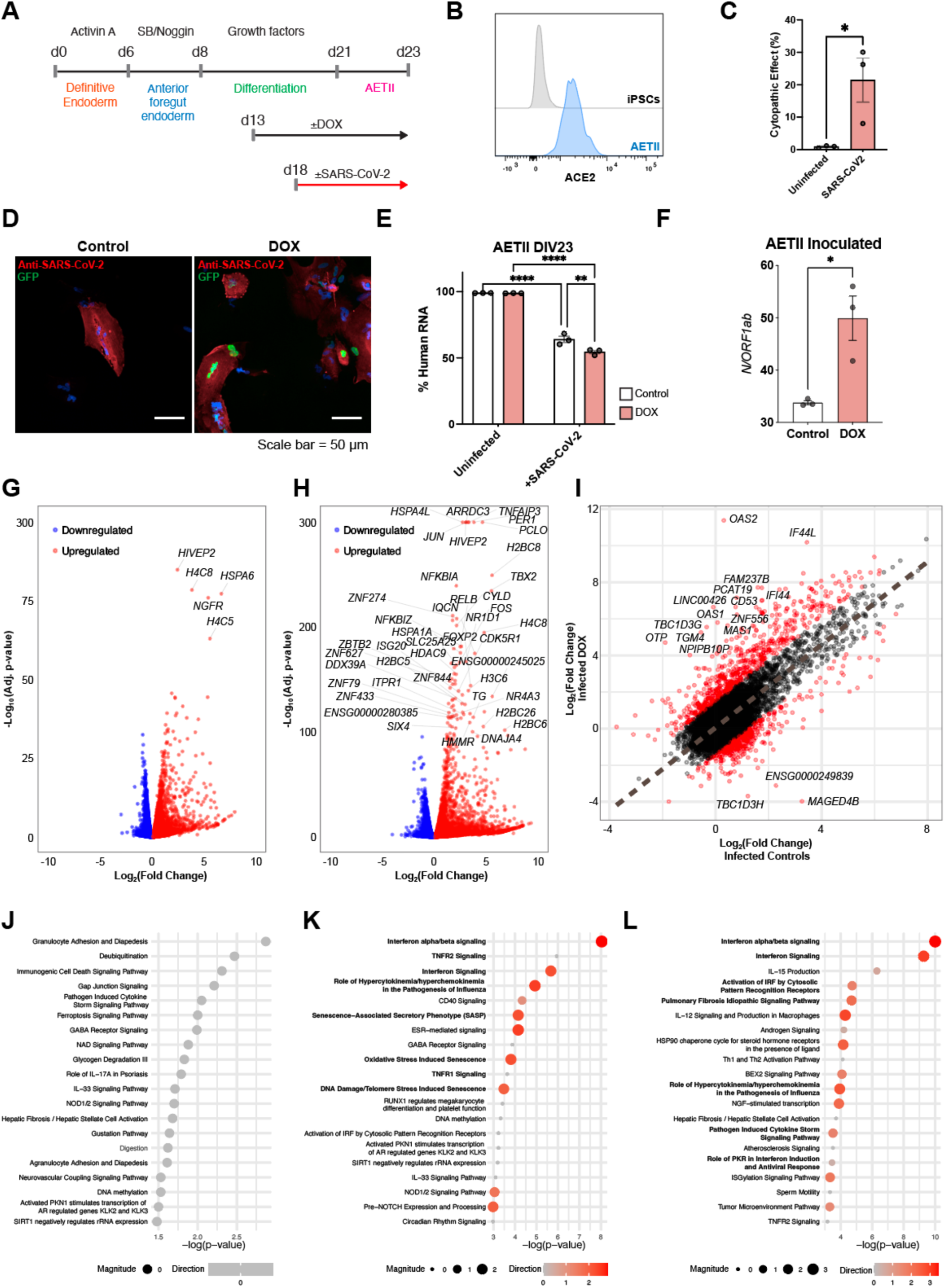
Progerin-induced AT2-like cells show enhanced inflammatory transcriptional responses to SARS-CoV-2. (A) Schematic of differentiation, induction and infection time points. (B) Flow cytometry showing ACE2 expression in iPSCs and AT2-like cells. (C,D) Cytopathic effect and SARS-CoV-2 antigen staining following exposure. N=3 experimental repeats. Statistical significance determined by Student’s t-test * P < 0.05. (E) Percentage of RNA-seq reads aligning to the human reference genome in differentiated-cell samples. Statistical significance determined by 2-Way ANOVA followed by Sydak’s test ** P < 0.01, **** P < 0.0001. N=3 experimental repeats. (F) RNA-seq quantification of SARS-CoV-2 transcript ratios in AT2-like epithelial cultures. Statistical significance was determined by Student T-test, * P<0.05. (G,H) Differentially expressed host genes after infection in uninduced and progerin-induced cultures. (I) Comparison of genome-wide infection-induced log₂ fold changes between progerin-induced and uninduced AT2-like cells. Each point represents one gene (n = 20,016). The dashed line represents an ordinary least-squares linear regression (R² = 0.63, slope = 1.134, intercept = 0.053). Red points indicate genes with absolute residuals above the 95th percentile of the fitted regression, highlighting transcriptional responses that deviate from the overall relationship. (J–L) IPA of infection responses and differences between infected states.

RNA-seq libraries from iPSCs showed a high proportion of reads aligning to the human reference genome irrespective of virus exposure (Supplementary Figure 2E), reflecting their inefficiency in supporting SARS-CoV-2 infection due to low expression of *ACE2* (Figure 2B, C, and 3B). By comparison, infected AT2-like cultures exhibited a reduced fraction of human-aligned reads, with a further decrease in infected progerin-induced samples (Fig. 3E). Consistent with this pattern, we found increased viral RNA (*N/ORF1ab*) in the DOX treated cultures supporting increased viral-genome loads (Fig. 3F).

Host transcriptional responses to infection differed in magnitude between the two cellular states. Volcano plots showed a broader and more pronounced pattern of differentially expressed host genes in progerin-induced cultures than in corresponding uninduced cultures (Fig. 3G,H). Genome-wide infection-associated changes were broadly correlated between conditions (Fig. 3I; ordinary least-squares linear regression, R² = 0.63, slope = 1.134), indicating that the overall transcriptional response to infection was largely shared between conditions. Nevertheless, a subset of genes showed a larger response with progerin induction, including interferon-stimulated genes such as *OAS1* and *OAS2*. Ageing- and senescence-associated interferon responses have also been described in other experimental systems [10].

Pathway analysis of the infection response in uninduced cells identified expected antiviral and innate immune program(Fig. 3J). In progerin-induced cells, interferon-associated pathways and inflammatory signatures were more prominent (Fig. 3K). Direct comparison of the infected conditions highlighted additional pathways annotated as hypercytokinaemia/hyperchemokinaemia, along with pulmonary fibrosis and cellular senescence-related signatures (Fig. 3L). These findings are consistent with amplified epithelial-intrinsic inflammatory signalling. Importantly, pathway names such as “cytokine storm” represent transcriptional associations; they do not demonstrate systemic hypercytokinaemia or increased secretion of inflammatory mediators in this monoculture model.

### Progerin induction is associated with increased infectious activity in culture supernatants

We next evaluated the relationship between progerin induction, antiviral treatment and the apparent production of infectious virus. Treatment with the Remdesivir metabolite GS-441524 markedly suppressed infection-associated cytopathic effects and viral-antigen staining (Fig. 4A,B). Although the progerin-induced group showed a directionally greater infection-associated signal in the untreated comparison, we did not detect any statistical differences specific to the ageing variable.

**Figure 4.**
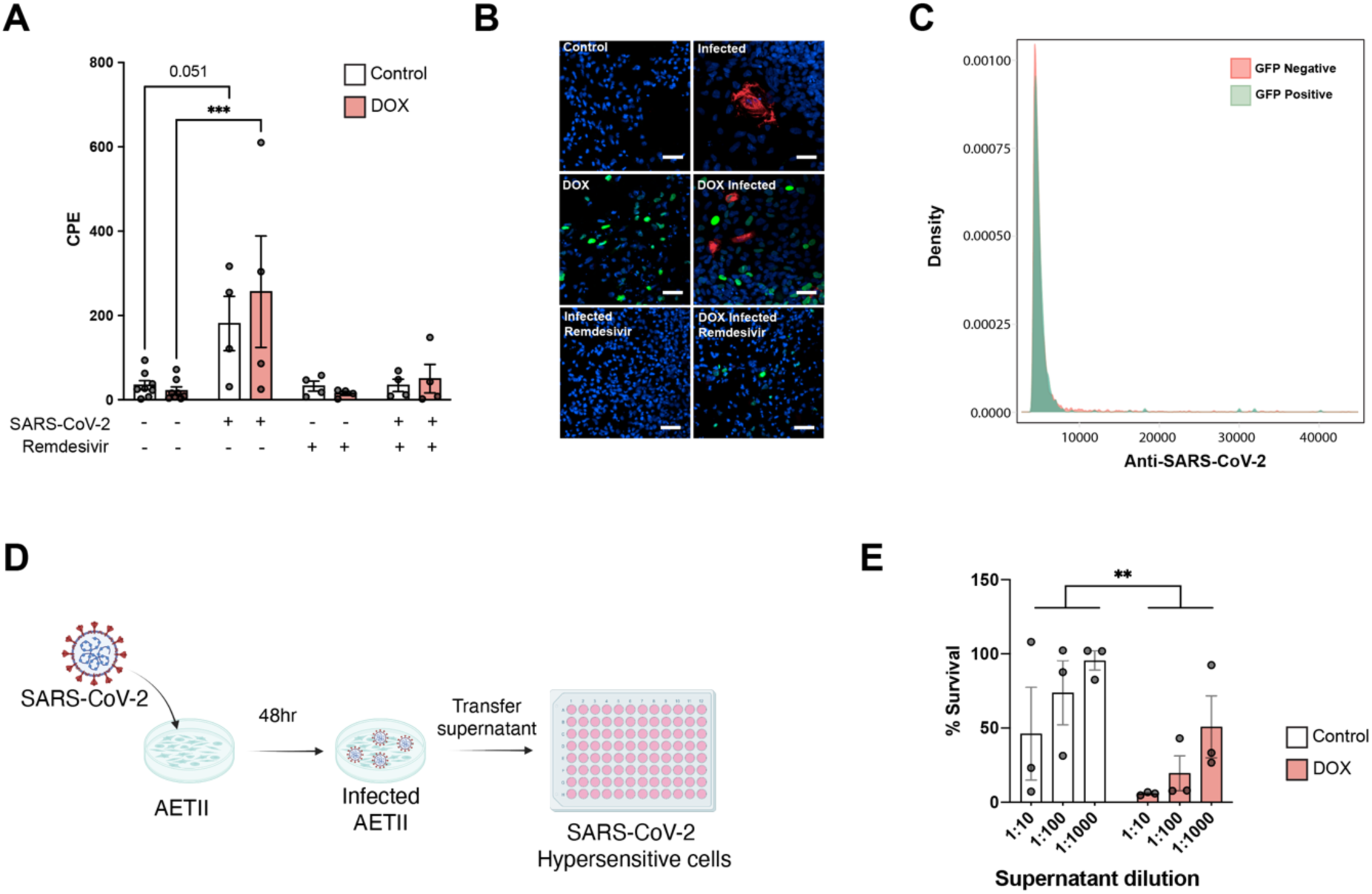
Antiviral response and supernatant-associated infectious activity in progerin-induced AT2-like cells. (A) Infection-associated readout with or without the antiviral intervention shown as remdesivir in the original panel. (B) Representative viral-antigen immunofluorescence images. (C) Distribution of SARS-CoV-2 antigen signal among progerin–GFP-positive and -negative cells. (D) Experimental scheme for transferring culture supernatants from infected AT2-like cells to a highly susceptible recipient-cell population. (E) Recipient-cell survival after exposure to serially diluted supernatants. Statistical significance was determined by 2-Way ANOVA.

Comparison of SARS-CoV-2 antigen staining between progerin–GFP-positive and GFP-negative cells did not reveal an obvious enrichment of infection in the GFP-positive fraction (Fig. 4C). Consistently, RNA-seq did not identify differential expression of the entry-related genes ACE2 or TMPRSS2 between induced and uninduced AT2-like cells (adjusted P = 0.70 and 0.88, respectively). These observations do not indicate a clear increase in receptor abundance or cell-level entry preference following progerin induction, supporting other innate mechanisms behind increased viral loads.

To obtain a functional measure of material released by infected epithelial cells, supernatants were collected after infection and transferred at serial dilutions to a SARS-CoV-2-hypersensitive reporter cell population (Fig. 4D). Supernatants from progerin-induced AT2-like cells contained significantly more viral particles leading to less survival in the recipient assay (Clonal HEK293 isolates expressing ACE2/TMPRSS2^10^) than those from uninduced controls at the tested dilutions (Fig. 4E).

## Discussion

This study establishes a human stem cell-derived epithelial model in which progerin can be introduced after early lineage specification to investigate the interaction between selected ageing-associated cellular states and SARS-CoV-2 infection. The system preserves an inducible iPSC platform, supports differentiation towards AT2-like cells and elicits changes in DNA-damage-associated staining, mitochondrial oxidant-associated fluorescence, heterochromatin marks and host gene expression. When challenged with SARS-CoV-2, progerin-induced cells display amplified inflammatory transcriptomic signatures and a functional supernatant phenotype consistent with increased infectious activity. The central contribution of our work is both methodological, a controllable route to introduce ageing-like features into differentiated cells, and biological, evidence for how the epithelial state established before exposure can modify the response to viral infection.

Pluripotent stem cell-derived respiratory models have provided important insight into epithelial susceptibility, virus-induced injury and candidate antiviral interventions^3,11^. Yet most models capture developmental or relatively youthful cell states^4^. AAVS1-integrated, doxycycline-regulated progerin expression creates a way to alter this context without requiring prolonged cell culture or introducing the ageing construct only after terminal differentiation. Such temporal separation is important because continuous progerin expression during stem-cell expansion could confound lineage specification. It also provides a platform for testing dose- and differentiation-stage-dependent effects, although these applications will require independent validation in additional cell types and genetic backgrounds.

The combination of elevated γH2AX, mitochondrial stress and reduced H3K9me3 provides convergent evidence that progerin changes several pathways associated with cellular ageing^6,7^. The transcriptomic induction of SERPINE1, CCN2 and inflammatory mediators further suggests a lung epithelial state that resembles selected features of tissue injury or age-related remodelling^12,13^. These phenotypes should not, however, be equated with physiological ageing. Progerin perturbs Lamin A biology and is associated with a rare premature-ageing disorder, whereas natural ageing reflects complex and heterogeneous processes^4,14^. The system is most accurately framed as a synthetic model of ageing-associated stress rather than a direct surrogate for an aged human lung.

The lower proportion of GFP-positive cells in differentiated cultures further suggests that the induced AT2-like population is heterogeneous with respect to detectable progerin–GFP expression (Supplementary Fig. 2A), this is likely a result of silencing in the differentiation process which can still affect the AAVS1 locus^15,16^. Future cell-resolved experiments will be necessary to assign these effects to individual expression states, while the exploration of other safe harbours e.g., CLYBL, ROSA26 could be tested^16^.

The transcriptional response to SARS-CoV-2 offers a possible connection between the pre-existing cellular state and inflammatory disease. Increased expression of OAS-family and interferon-responsive genes is consistent with a stronger antiviral host response^3,11^, while enriched profibrotic and cytokine-related signatures suggest that the same state may amplify programs associated with tissue injury^11^. The magnitude of an antiviral transcriptional response does not necessarily reflect its effectiveness in restricting infection, as the timing of interferon-stimulated gene induction relative to viral replication can critically influence antiviral protection^17,18^. Indeed, delayed interferon responses have been observed following SARS-CoV-2 replication in human respiratory epithelial models^19^, while pre-existing cellular stress may independently contribute to inflammatory signalling. Previous studies have demonstrated that progerin-induced replication stress and genomic instability can activate STAT1-regulated interferon-like responses^20,21^, suggesting a potential intersection between ageing-associated nuclear stress and antiviral signalling. Thus, the enhanced transcriptional response observed in our model could reflect increased viral burden, progerin-associated inflammatory priming, or a combination of these processes.

Our results support future studies to model “cytokine storm” features that are associated with the aged population. By incorporating epithelial, stromal, vascular and immune compartments subsequent work can characterised innate and secreted mediators of this clinical feature^2,11^. To elaborate, future co-culture or multicellular organoid systems, together with secretome profiling, would help examine whether the epithelial signals identified here propagate inflammatory responses to neighbouring cell populations.

The infection-related findings further support an altered host–virus interaction in progerin-induced AT2-like cells. The reduced proportion of sequencing reads aligning to the human genome was accompanied by increased SARS-CoV-2 RNA abundance, as determined by quantification of viral *N/ORF1ab* transcripts. Consistent with these observations, supernatants collected from progerin-induced cultures produced greater cytopathic activity in highly susceptible recipient cells, supporting increased infectious activity in the conditioned medium (Fig. 4D,E). Interestingly, these findings occurred alongside enhanced interferon-associated and inflammatory transcriptional responses, suggesting that heightened antiviral signalling does not necessarily translate into more effective restriction of infection. These findings are consistent with progerin-associated cellular stress altering the interaction between SARS-CoV-2 and the host epithelial response, resulting in increased viral RNA abundance and infectious activity despite amplified inflammatory signalling.

In summary, inducible progerin expression enables the study of selected ageing-associated phenotypes in human stem cell-derived alveolar epithelial-like cells and reveals a heightened inflammatory response to SARS-CoV-2 infection. By making cellular state an experimentally adjustable variable, this platform may support investigations of respiratory infection and other disease processes in which age-associated epithelial biology is relevant.

## Materials and methods

### Human iPSC maintenance and inducible cassette engineering

The human iPSC line WTC-C ^22^ was used as the parental and control line. Related lines are further described here ^23^. Cells were maintained under feeder-free conditions in mTeSR1 medium (STEMCELL Technologies) on Matrigel-coated plates (354277, Corning). A ROCK inhibitor (Y-27632, 10 µM) was supplied transiently for less than 24 h following cell passage. An inducible cassette was derived from a previously reported AAVS1-targeted strategy ^5^ by replacing the original Cas9 coding region with a progerin–GFP insert^24^. AAVS1 targeting was performed using TALEN-mediated integration as originally described, with puromycin- and neomycin-resistance selection cassettes depicted in Fig. 1A^5^.

### Differentiation towards alveolar type II-like cells

Cells were differentiated using a multistage lung epithelial protocol (Fig. 2A)^25^. Briefly, definitive endoderm differentiation was initiated by culturing iPSCs in RPMI 1640 medium supplemented with Activin A (100 ng/mL), L-glutamine (2 mM) for 48 h. Cells were subsequently cultured for an additional four days in the same medium supplemented with B27 (1X), sodium butyrate (0.5 mM), and fetal bovine serum (FBS; 0.1%), with daily medium changes. On day 6, definitive endoderm cells were dissociated and replated onto matrigel-coated plates for anterior foregut endoderm specification. Cells were cultured for 48 h in Iscove’s Modified Dulbecco’s Medium (IMDM) supplemented with 5% FBS, 2 mM L-glutamine, 1 mM non-essential amino acids, Noggin (200 ng/mL), and SB431542 (10μM). From day 8, cells were directed towards an alveolar epithelial lineage using IMDM supplemented with 10% FBS, 2 mM L-glutamine, 1 mM non-essential amino acids, retinoic acid (0.5 µM), fibroblast growth factor 10 (FGF10; 10 ng/mL), epidermal growth factor (EGF; 10 ng/mL), Wnt3a (100 ng/mL), and keratinocyte growth factor (KGF/FGF7; 10 ng/mL). Differentiation was continued until approximately days 21–23 to generate cells with an alveolar epithelial phenotype. To induce Progerin expression, doxycycline (DOX; 1 µg/mL) was added to the culture medium beginning on day 13 and maintained throughout the subsequent experimental interval. Parallel cultures maintained without doxycycline served as controls.

### Quantitative RT-PCR and lineage characterisation

RNA was isolated from the indicated iPSC and differentiated-cell populations, and transcript abundance was measured by reverse-transcription quantitative PCR using the ΔΔCt method for the lineage and viral-entry markers. A full list of primers is attached in Supplemental Table 1.

### Immunofluorescence and microscopy

Cells were fixed in 4% paraformaldehyde for 20 min at room temperature, permeabilised with 0.3% Triton X-100, and blocked for 1 h in PBS containing 0.05% Triton X-100, 5% normal goat serum and 1% bovine serum albumin, as described in the original draft. Primary antibodies were incubated overnight at 4 °C in blocking solution. Samples were washed, incubated with fluorescent secondary antibodies for 1 h at room temperature, washed again and imaged. For a full list of antibodies used in this study see Table 1.

High-content images were acquired using an Opera Phenix platform (PerkinElmer) and analysed in Harmony software. Additional images were collected with an IN Cell Analyzer and analysed using CellProfiler. DNA-damage-associated staining was quantified using nuclear γH2AX signal or foci. Nuclear H3K9me3 was assessed using fluorescence intensity distributions measured around in the nucleus as defined by masking with hoechst staining.

### Mitochondrial oxidative stress and senescence-associated β-galactosidase

Mitochondrial oxidant-associated fluorescence was assessed by live-cell MitoSOX staining (M36007, ThermoFisher Scientific), and signal intensity was compared among induced and uninduced conditions and by GFP status. Senescence-associated β-galactosidase activity was assessed in parallel (#9860, Cell Signaling Technologies)

### Flow cytometry

Single-cell suspensions were prepared using Accutase and resuspended in flow cytometry buffer consisting of PBS and 1% bovine serum albumin. For ACE2 staining, the original draft reports a 20 min blocking step at room temperature, primary antibody incubation at a 1:400 dilution for 20 min, two washes, and incubation with a donkey anti-goat Alexa Fluor 633 secondary antibody at 1:500 for 20 min. Samples were analysed using a Cytek Aurora 3L instrument.

### SARS-CoV-2 infection and antiviral response analysis

Experiments involving infectious SARS-CoV-2 were conducted under biosafety level 3 containment. The study used the Ancestral SARS -CoV-2, with virus stocks and infection procedures as referenced ^10,26^. Cells were analysed for infection-associated cytopathic effects and viral-antigen staining at the indicated collection time. An antiviral comparator described in the figure as remdesivir was applied to infected cultures at a reported concentration of 5 µM GS-441524 (72-271-0, Thermofisher Scientific).

### Supernatant-transfer infectivity assay

Culture supernatants were collected from SARS-CoV-2-exposed AT2-like cells after 48 hours of inoculation, and serially diluted aliquots were transferred to a highly susceptible recipient-cell assay, as described in ^10^. Recipient-cell survival was determined 48 hours post-inoculation using standardized automatic image quantifications as previously ^10,26^.

### RNA sequencing and differential-expression analysis

Bulk RNA sequencing was performed on the indicated iPSCs, differentiated, progerin-induced and SARS-CoV-2-exposed samples collected from 3 independent experiments. Sequencing was performed by the Ramaciotti Centre for Genomics using the NovaSeq2000 platform. Sequencing reads were aligned to a human reference genome and summarised to quantify gene-level expression. The fraction of reads aligning to the human genome was compared across conditions, separately human and SARS-CoV-2 genome alignment was performed to quantify viral expression.

Differential gene expression analysis was performed using DESeq2 in R. Raw gene counts were normalised using DESeq2 size-factor estimation, and genes were retained if they exhibited at least 10 normalised counts in two or more samples. Differential expression was assessed using a negative binomial generalised linear model, with experimental condition included as the design factor. Statistical significance was evaluated using the Wald test, and P values were adjusted for multiple comparisons using the Benjamini–Hochberg false-discovery rate procedure. For stringent selection of differentially expressed genes, thresholds of adjusted P < 0.01, absolute log₂ fold change > 3, and mean normalised counts > 400 were applied. Gene-level results were visualised using volcano plots displaying log₂ fold changes against −log₁₀-adjusted P values. Genes with absolute log₂ fold changes > 0.6 and adjusted P < 0.001 were selected for annotation. Heatmaps were generated from size-factor-normalised expression counts, transformed to gene-wise Z-scores to visualise relative expression differences across samples. Selected lung epithelial, developmental and pluripotency markers were visualised using curated gene lists.

Ingenuity Pathway Analysis (QIAGEN) was used to evaluate enrichment and predicted activation or inhibition of curated biological pathways. Pathway enrichment was assessed using the software’s statistical framework, while predicted activation states were evaluated using activation Z-scores where available.

### Statistical analysis

Statistical analyses were conducted using Prism 11.1.0 (GraphPad Software Inc.) unless indicated. N indicates the number of independent experiments. Data are represented as the mean ± SEM where applicable. When relevant, all statistical analyses were corrected for multiple comparisons. Additional information about statistical tests is reported in the figure legends.

## Data and code availability

The RNA-sequencing datasets generated in this study have been deposited in the National Center for Biotechnology Information (NCBI) Sequence Read Archive (SRA) under BioProject accession number PRJNA1560203. The datasets are currently held under embargo and are scheduled for public release on 1 December 2027.

## Acknowledgements

GGN was supported by the NHMRC (GNT2020532, GNT1185002, GNT1107514, GNT1158164, GNT1158165, GNT1046090, GNT1111940) a kind donation from Dr. John and Anne Chong. CLM was supported by a kind donation from Dr. John and Anne Chong. The authors would like to acknowledge the Sydney Cytometry Facility and the Sydney Microscopy Facility for technical assistance.

## Author contributions

CLM acquired and analyzed data (SARS-CoV-2 in vitro infections, stem cell characterizations and generation), and drafted the manuscript. AA contributed by performing viral inoculations and scientific planning. ST helped in scientific planning and differentiation. DV participated in the generation and characterization of stem cell lines. ST, JCW, EW, GGN helped in the general planning of experiments, project management, and manuscript edits.

## Ethics and biosafety approvals

All lines used in these studies were approved by the University of Sydney Human Ethics Committee (HREC 2020/417).

## Supplementary Figure Legends

**Supplementary Figure 1.**
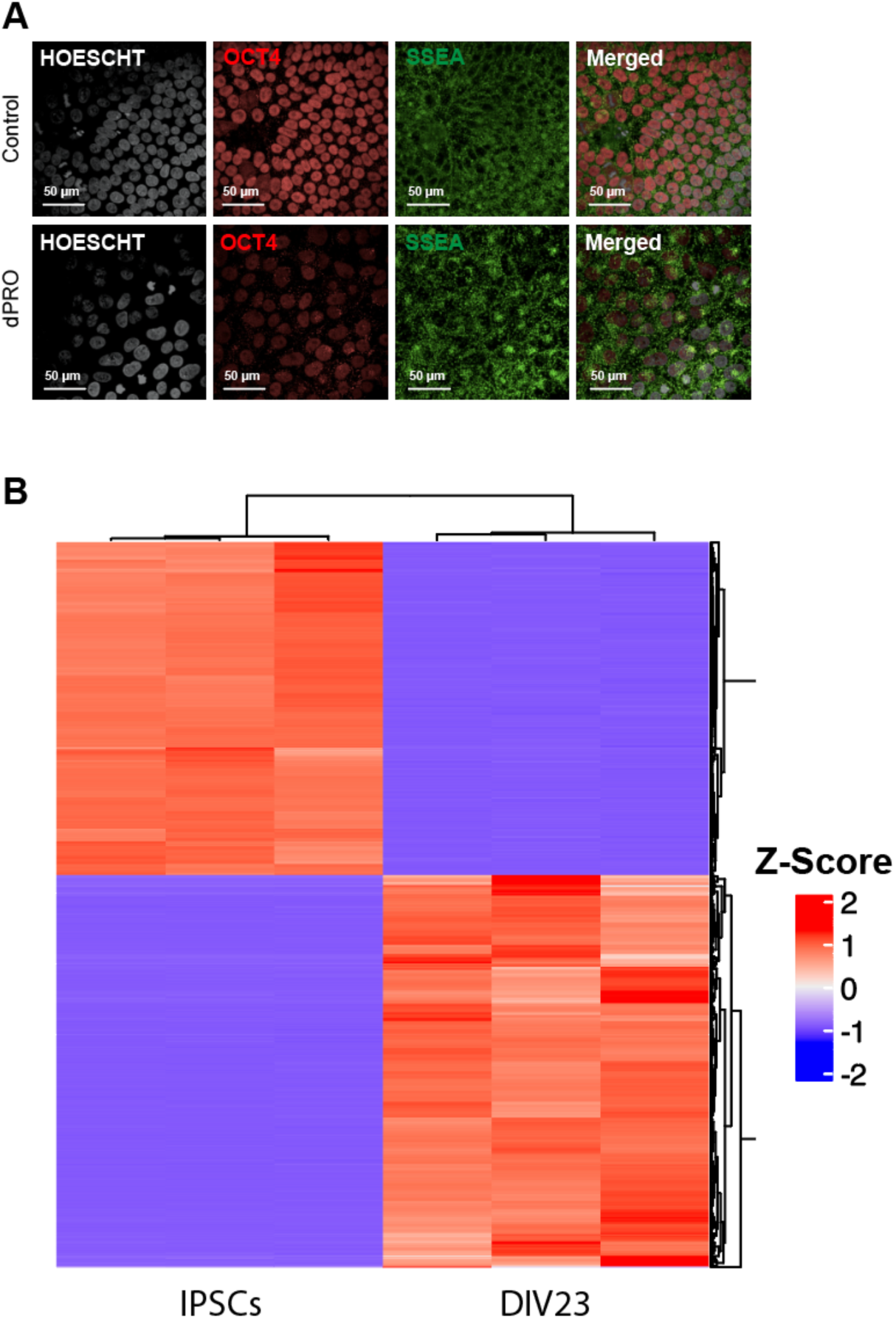
Pluripotency-marker expression and transcriptomic divergence during AT2-like differentiation. (A) Representative immunofluorescence images of parental control and inducible progerin (dPRO) iPSCs stained for OCT4 (red) and the SSEA-4 pluripotency-associated surface marker (green), with Hoechst nuclear staining (grey) and merged images. Scale bars, 50 µm. (B) Heatmap comparing the expression profiles of undifferentiated iPSCs and day-23 differentiated AT2-like cultures (DIV23). Rows represent transcripts and columns represent individual samples; gene-wise Z-scores indicate relative expression (blue, lower; red, higher). Hierarchical clustering separates the two culture states.

**Supplementary Figure 2.**
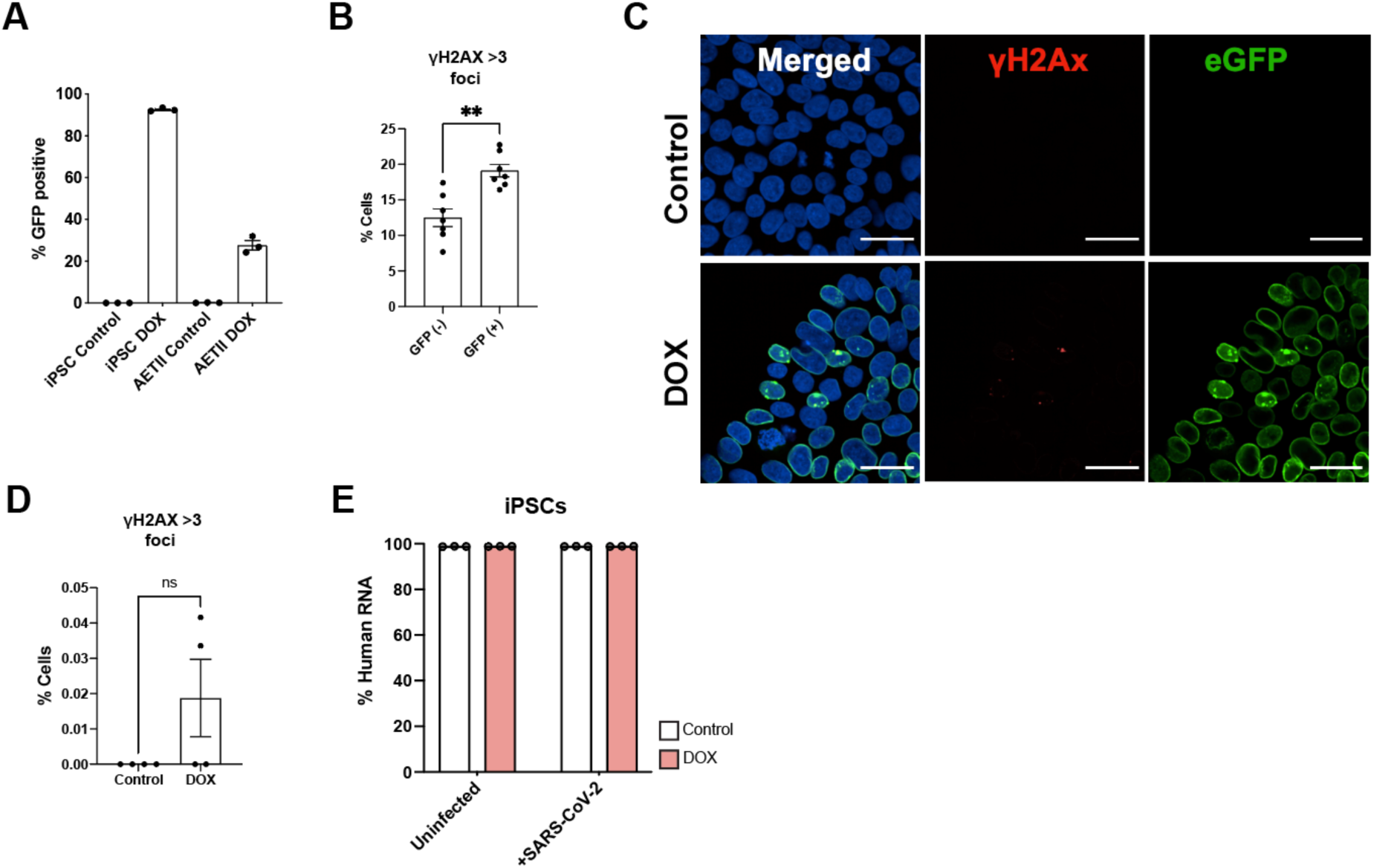
Differentiation-dependent progerin induction and γH2AX-associated DNA damage. (A) Percentage of GFP-positive cells in untreated control and doxycycline (DOX)-treated iPSCs and AT2-like epithelial cultures, showing a lower GFP-positive fraction in differentiated cultures following induction. (B) Percentage of GFP-negative and GFP-positive AT2-like cells containing more than three nuclear γH2AX foci. The displayed comparison is significant (**P < 0.01). (C) Representative microscopy of untreated and DOX-treated iPSCs stained for γH2AX (red), progerin-associated GFP (green) and nuclei (Hoechst, blue); merged images are shown at left. (D) Quantification of the frequency of iPSCs with more than three γH2AX nuclear foci in untreated and DOX-treated conditions; ns, not statistically significant. Symbols represent individual experiments. (E) Percentage of RNA-seq reads aligning to the human reference genome in iPSC N=3 experimental repeats. Statistical significance determined by 2-Way ANOVA followed by Sydak’s test ** P < 0.01, **** P < 0.0001.

