## Supplemental Tables for "Progerin Expression in Alveolar Epithelial Cells Exacerbates Inflammatory Responses to SARS-CoV-2 Infection"

### Supplementary Materials

### Supplemental Table 1. Oligonucleotides

**Primers for correct gene targeting**

| **Primer name** | **Sequence** |
|---|---|
| DNA803 | TCGACTTCCCCTCTTCCGATG |
| DNA183 | CTCAGGTTCTGGGAGAGGGTAG |
| DNA804 | GAGCCTAGGGCCGGGATTCTC |
| DNA458(PURO) | CTCGACATCGGCAAGGTGTG |
| DNA1131(NEO) | TGAATGAACTGCAGGACGA |
| PURO_R | GAGTTCTTGCAGCTCGGTGA |
| NEO_R | AATGGCCGATCCCATACGC |

See reference for assessing correct gene targeting
https://doi.org/10.1016/j.ymeth.2015.12.012

RT-PCR primers

| **Name** | **Sequence** |
|---|---|
| *NANOG F* | CCAAATTCTCCTGCCAGTGAC |
| *NANOG R* | CACGTGGTTTCCAAACAAGAAA |
| *NKX2.1 F* | TAACGGCGGCAGACAAAAAGA |
| *NKX2.1 R* | GAAGTATTGCTTCAGTTGGCCT |
| *ACE2 F* | GGACCCAGGAAATGTTCAGA |
| *ACE2 R* | GGCTGCAGAAAGTGACATGA |
| *SFTPC F* | CCTTCTTATCGTGGTGGTGGT |
| *SFTPC R* | TCTCCGTGTGTTTCTGGCTCAT |
| *GAPDH F* | acaactttggtatcgtggaagg |
| *GAPDH R* | gccatcacgccacagtttc |
| *MUCIN F* | AGCTTCTACTCTGGTGCACAA |
| *MUCIN R* | GGTGGCTGGGAATTGAGA |

**Supplemental Table 2. Antibodies**

### Primary Antibodies

| Target | Manufacturer | Catalog | Dilution |
|---|---|---|---|
| ACE2 | Thermofisher | MA532307 | 1 in 200 (immunocytochemistry) |
| ACE2 | R&D Systems | AF933 | 1 in 400 (flow cytometry) |
| gammaH2AX | Cell Signalling Technologies | 9718T | 1 in 400 |
| H3K9-Trimethylated | abcam | ab176916 | 1 in 500 |
| NKX2.1 | abcam | ab76013 | 1 in 200 |
| Oct4 | Thermofisher | A24867 | 1 in 200 |
| Podoplanin | Thermofisher | 14-9381-80 | |
| SARS-CoV-2 Serum | Made in house (REF) | NA | NA |
| SOX2 | Thermofisher | A24759 | 1 in 100 |
| SSEA4 | Thermofisher | A24866 | 1 in 100 |
| TMPRSS2 | Thermofisher | PA5-14264 | 1 in 100 |
| TRA-1-60 | Thermofisher | A24868 | 1 in 100 |

### Secondary Antibodies

| Target | Manufacturer | Catalog | Dilution |
|---|---|---|---|
| Rabbit IgG 647 | Thermofisher | A21244 | 1 in 1000 |
| Rabbit IgG 555 | Thermofisher | A21428 | 1 in 1000 |
| Rat IgG 647 | Abcam | Ab150167 | 1 in 1000 |
| Mouse IgG 555 | Thermofisher | A21422 | 1 in 1000 |
| Human IgG 647 | Thermofisher | A21445 | 1 in 1000 |
